# A Framework for Sensor-based Assessment of Upper-limb Functioning

**DOI:** 10.1101/2021.02.10.430700

**Authors:** Ann David, Tanya Subash, SKM Varadhan, Alejandro Melendez-Calderon, Sivakumar Balasubramanian

## Abstract

The ultimate goal of any upper-limb neurorehabilitation procedure is to improve upper-limb functioning in daily life. While clinic-based assessments provide an assessment of what a patient can do, they do not completely reflect what a patient does in his/her daily life. The compensatory use of the less affected upper-limb (e.g. “learned non-use”) in daily life is a common behavioral pattern seen in patients with hemiparesis. To this end, there has been an increasing interest in the use of wearable sensors to objectively assess upper-limb functioning. This paper presents a framework for assessing upper-limb functioning using sensors by providing: (a) a set of definitions of important construct associated with upper-limb functioning; (b) presenting different visualization methods for evaluating upper-limb functioning, along ways to qualitatively analyze different visualization methods; and (c) two new measures for quantifying how much an upper-limb is used and the relative bias in the use of the two upper-limbs. The demonstration of some of these components is presented using data collected from inertial measurement units from a previous study. The proposed framework can help guide the future technical and clinical work in this area to realize a valid, objective, and robust tool for assessing upper-limb functioning. This will in turn drive the refinement and standardization of the assessment of upper-limb functioning.

## 1 Introduction

After neurological injury, individuals require rehabilitation to promote recovery, minimise disability and maximise independent living. Despite years of research pointing to the benefits of repetitive practice, the time patients spend in inpatient rehabilitation settings is often much less than the recommended guidelines [1, 2]. Moreover, after discharge, patients do not have enough opportunities to do targeted movement therapy at home, sometimes leading to a pattern of “learned non-use” [3].

Valid and reliable assessments are crucial for gaining a better understanding of impairments and recovery potential, and can allow us to tailor intervention strategies or improve health services. While clinic-based assessments of body function and activity can measure the capability of a patient, they are poor indicators of the actual use of a limb in day-to-day life [4–6]. Thus, assessment of movement behavior in natural settings is vital to evaluate recovery and the tangible impact of rehabilitation interventions. In the context of hemiparesis, such assessments can help gauge the extent to which: (a) true recovery in the more-affected limb is utilized for daily use, and (b) compensatory strategies with the less-affected limb and other body parts is used to accomplish day-to-day activities. To this end, assessments such as the motor activity log (MAL) [7] have been devised to capture upper-limb functioning of patients with hemiparesis.

There are four inter-related aspects that need consideration to depict a comprehensive picture of upper-limb functioning:

- **Question 1**. How much is an upper-limb used during daily life?
- **Question 2**. What is the relative preference for using the more-affected limb over the less-affected one?
- **Question 3**. What kind of tasks is an upper-limb is used for?
- **Question 4**. What is the quality of upper-limb movements?

The first two questions convey information about how much the upper-limbs are used and their relative preference, the third question provides information about the nature of use of the upper-limb, and the last question gives insights about the underlying motor control abilities. The MAL is structured to gather this information [7] where the amount and quality of use are rated on a 6-point Likert scale for a set of pre-selected tasks. The amount of use of the more-affected limb is reported by comparing it to amount of use of the less-affected limb. The quality of use of the less-affected limb is reported with respect to the pre-stroke condition of that limb. However, MAL can only provide a coarse and subjective evaluation of upper-limb functioning in daily life due to its limited sensitivity and subjective nature as it relies on a patient’s ability to recall upper-limb use from memory.

There is growing interest in wearable sensors for continuous and objective monitoring of upper-limb functioning [8–16]. When developing a sensor-based assessment tool, there are a four major interdependent design choices that influence the nature of the information conveyed by the assessment: (a) the type of sensing modality used for measurements (e.g., camera-based movement tracking, inertial measurement units), (b) steps involved in the data processing pipeline (e.g., data segmentation, filtering), (c) properties of measures used to quantify constructs of interest (e.g., sensitivity to movement changes in the physiological changes, robustness to measurement noise), and (d) the nature of data visualization methods employed (e.g., temporal evolution of the measure, scatter plots of different variables). Some of these issues have been highlighted previously for analysing movement smoothness [17–19].

Inertial sensors composed of accelerometers and gyroscopes have been the preferred modality for assessing upper-limb functioning in the natural setting, due to their availability, affordability, and compact size [8–16]. Thus far, the focus of sensor-based assessment in hemiparesis has been the quantification of the overall amount (question 1) and the relative bias (question 2) in using the arms during daily life [8–16]. The current methods for quantifying the amount of upper-limb use have either used: (a) the magnitude of acceleration (e.g. activity counting (AC) [8, 11, 20]) or (b) the duration of functional movements detected from sensor data (e.g. gross movement (GM) score [10, 13], machine learning (ML) algorithms [14–16]). Although related, movement duration and intensity convey slightly different information about the nature of arm use. Each of these only provide partial characterization of how much a particular arm is used. A complete measure of how much an arm is used in daily life requires knowledge of both the duration and the intensity of the upper-limb movements. Also, there is currently little work using sensor data for determining the nature of tasks/activities and quantifying the quality of movements performed. These aspects are likely to be explored in the coming years with the increasing interest in this area, the availability of more data and sophisticated data analysis methods.

In order to develop rigorous methods to assess different aspects of upper-limb functioning in daily life, now is an opportune moment to lay a good foundation for this problem through a more formal framework consisting of: (a) definitions of essential concepts, and (b) recommendations for the development, and analysis of quantitative measures and (c) visualization methods. Such a framework can help steer future technical developments in the appropriate direction, and limit work on ill-founded methods and procedures.

This paper presents a framework for sensor-based upper-limb functioning assessment, targeting researchers developing and validating quantitative methods. Given the multi-disciplinary nature of the community, our goal is to unify the language with formal definitions, and have attempted to convey the core ideas with as little mathematical formalism as possible. The framework presented in this paper starts with formal definitions of the relevant concepts in assessment of upper-limb functioning related to aforementioned questions (Section 2). This is followed by different approaches for visualizing quantified constructs, such as how much an upper-limb is used, and the relative bias between the two upper-limbs (Section 3). Additionally, methods for qualitative understanding of the nature of a visualization approach and its interpretation are also presented. The proposed visualization methods are also accompanied by quantitative measures that summarize nature of distribution of data in these methods. The rationale for these measures and the qualitative validation of their properties are also provided. We conclude the paper with a discussion of the limitations of the proposed framework, along with important questions that must be addressed to make pervasive, sensor-based objective assessment of upper-limb functioning in daily-life a clinical reality.

## 2 Measuring Upper-limb Functioning: Formal definitions

We use our upper-limbs for performing movements and postures [21] for accomplishing various tasks in our daily life. The nature, amount, intensity, and quality of movements performed with the upper-limbs are determined by: (a) the types of tasks performed; (b) the overall motor ability; and (c) the hand preference. This is depicted in Fig. 1. While motor ability can be evaluated through standard clinical assessments employed in the clinic (e.g., FMA, ARAT), assessments of upper-limb functioning in natural settings is necessary to evaluate how much and how well the upper-limbs are incorporated in daily life activities. This can be gauged by investigating the following inter-related aspects:

- Amount of functional versus non-functional use of the upper-limbs.
- Total duration and intensity of use.
- Relative use of one upper-limb over the other.
- Amount of uni- versus bi-manual use.
- Types of tasks performed.
- Quality of movements.

**Figure 1:**
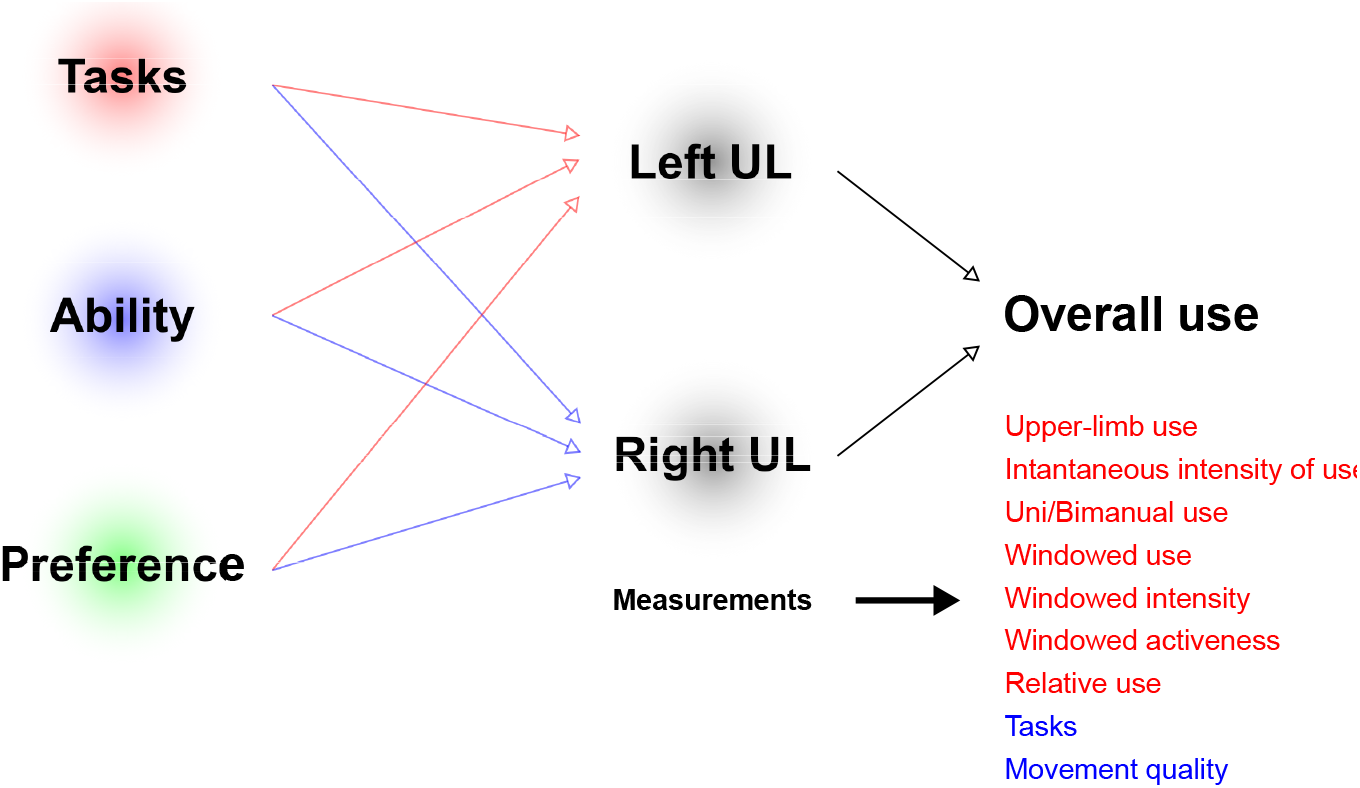
Different factors affecting the nature of use of the two upper-limbs during daily life. The type of task, the ability of a subject to perform it, and his/her preference for the particular arm to complete the task affect the nature and amount of upper limb movements. Sensors record these movements and assess the overall upper-limb use through quantification of different constructs.

The quantification of these different aspects can help us distinguish between a patient’s true recovery versus compensation [22]. To promote the development of appropriate methods to quantify these different aspects, we require clear definitions for these aspects, which is the purpose of this section.

Before getting into the details of the framework, we start with a brief overview of the process of evaluating the sensorimotor ability/function in the context of neurorehabilitation. This is a hierarchical process with clinical evaluation at its highest level. We define an **evaluation** as the process of interpreting the results of one or more assessments to gauge the sensorimotor condition of a subject with respect to a reference (either him/herself from a different time point (intra-subject), or another subject (inter-subject)). For instance, an evaluation is performed when comparing the results of ARAT assessments across different time points, or comparing smoothness of reaching movements of a patients against normative data. Evaluations can be aided through visualizations that allow interpretation of assessments. The next level in this hierarchy are **assessments**, which we define as the process of quantifying (i.e. putting numbers) abstract theoretical constructs (e.g., smoothness, coordination, synergies). For instance, the Fugl-Meyer upper-limb assessment is a process that aims at quantifying the constructs ‘motor function’, ‘synergy’ and ‘coordination’. Unlike an evaluation, an assessment only deals with quantifying constructs of interest. Assessments require clearly defined protocols for collecting data (e.g. tasks/movements to be performed), and measures. A **measure** is a well-defined mathematical function/formula, a computational algorithm, or a set of processes for mapping measurements or observations to quantities that have an interpretable meaning in the context of a construct. For instance, SPARC and LDLJ are measures of the construct ‘movement smoothness’; the rules used for assigning a score to the flexion synergy task in the Fugl-Meyer assessment is a measure of the construct ‘flexion synergy’. Measures with good properties are essential to obtain valid, reliable, and interpretable assessments. Finally, **measurements** are records of variables (e.g.speed, position, orientation, etc.), which could be obtained through various sensors or through human observation. In this section we define some of the important terms and constructs associated with the assessment of upper-limb functioning.

### 2.1 Measurement space

Raw measurements of movement-related variables using sensors form the basis of all sensor-based assessments. The type of measurements available determine the subsequent steps in the analysis process. It is thus crucial for any assessment procedure to clearly state the variables that are being used to compute a specific measure. In the context of the assessing upper-limb functioning, we define **measurement space** as the following.

*Definition* **Measurement space** is the universal set of all possible sensor measurements available from an upper-limb that is input to a measure.

We denote this set – the measurement space – by 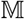 and assume that the same quantities are measured from both upper-limbs for the given assessment procedure. Inertial sensing is one of the most common modalities used in the current literature, where the arm movements are measured using wrist-worn accelerometers 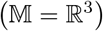 or IMUs^2^ 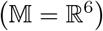. For a more elaborate measurement setup consisting of measurements of wrist endpoint position and orientation, along with *k* joint angles, 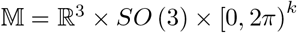; where ℝ^3^ is the set of all possible wrist positions, *SO* (3) is the special orthogonal group of all rotation matrices representing 3D orientations of the wrist, and [0, 2*π*)^*k*^ is the set of *k* joint angles.

All measures use measurements made over a finite observation period referred to as the *measurement epoch*. Let *M*_*l*_ (*t*) and *M*_*r*_ (*t*) represent the values of the measurements from the left and right upper-limb, respectively, made at time instant *t*, where *t* ∈ [0*, T*], and *M*_*l*_ (*t*), 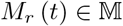; *T* is the duration of the measurement epoch. We use *M*_*l*_ and *M*_*r*_ to represent the entire time series or signal, where *M*_*l*_, 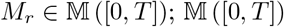 is the set of all possible measurement signals over a measurement epoch of duration *T* seconds starting at time *t* = 0. In addition to specifying 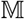, it is also essential for the reproducibility of an assessment to clearly specify the exact sensors used for the measurements, their accuracy, noise characteristics, resolution, sampling rate, etc. The values of these parameters have practical implications during data analysis and interpretation. These practical issues will be not be considered in this manuscript, and the mathematical formalism is presented assuming that we are dealing with measurements that are continuous in time and space.

### 2.2 Upper-limb use

*Definition* **Upper-limb use** is a binary construct indicating the presence or absence of voluntary, meaningful movement or posture of the limb.

In this definition, the boundary of what constitutes a “meaningful” movement/posture must be defined apriori. Some examples of meaningful use include reaching and grasping, turning a doorknob, stabilizing an object with one limb while manipulating it with the other, holding a glass, writing, typing, upper-limb therapy exercises etc. Under this definition, involuntary and passive upper-limb movements/postures are not considered meaningful, e.g., resting the arm on a table, upper-limb moved by an external force. There are, however, cases where the presence/absence of upper-limb use is ambiguous, e.g., arm swing during walking, passively resting the upper-limb on a book to prevent the pages from turning. Such ambiguities are best resolved in an application-specific manner, where the set of tasks considered as meaningful are clearly stated apriori. For instance, in the current upper-limb use literature arm swing during walking is not considered as meaningful, even though these are unlikely to be purely passive movements [23].

Upper-limb use can be mathematically represented as a binary signal over time, which can be computed from upper-limb measurements 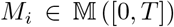, where *i* ∈ {*l, r*}. Let *f*_*u*_ be a function representing a measure that maps a given measurement signal *M*_*i*_ to a binary signal *u*_*i*_ over the same temporal domain, i.e. 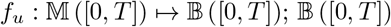 is set of Riemann-integrable binary signals in the time interval [0*, T*].

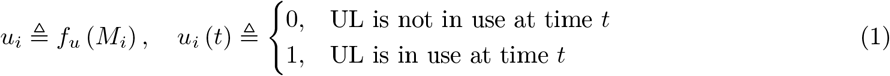

where, *u*_*i*_ is the upper-limb use signal of the upper-limb *i*. The choice of *f*_*u*_ is determined by several factors, e.g., the measurement space 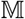, computational complexity of the measure, sensitivity/specificity of the algorithm. All *f*_*u*_ exploit some common structure in functional/meaningful movements present in the measured data to detect upper-limb use. Some examples of the current methods (*f*_*u*_) are:

- **Thresholded activity counting**. Activity counting (AC) is one of the most popular methods in the literature to quantify upper-limb functional and non-functional activity [8, 9, 11, 20]. AC has high sensitivity, but poor specificity [24]. AC can be used with both accelerometer and IMUs. Upper-limb use *u*_*i*_ is computed from AC by assigning a value of 1, whenever the AC is above a threshold.
- **Gross movement (GM) score**. The Gross movement (GM) score (a.k.a Gross Counts or Gross Movement Identification method) proposed by Leuenberger et. al [13] reconstructs the forearm orientation using a wrist-worn IMU to detect movements that occur in a pre-specified range of forearm orientations [13]. The GM score is highly specific, but has low sensitivity [24]. The GM score can only be used with an IMU. The GM score is 1 whenever there are arm movements in a pre-specified range of forearm orientations.
- **Random Forests classifier**. Bochniewicz et. al [15] proposed the use of a random forests classifier to detect upper-limb use from features extracted from an accelerometer. The ML approach can be used with both accelerometers and IMUs, and has reasonable sensitivity and specificity [16].

The three aforementioned methods are possible measures of upper-limb use and one must be aware of the properties of these measures while choosing them for the assessment of upper-limb use.

Upper-limb use as defined in this section is an idealised construct, and its detection in practice using sensor measurements will be error prone due to measurement noise, the natural intra- and inter-subject movement variability, and the relative sensitivity of the sensor measurements to movements and postures. The nature of the measurements 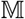 and the choice of measure *f*_*u*_ will influence how well upper-limb use can be quantified (e.g., sensitivity and specificity) in practice. For instance, among the three measures currently used in the literature, AC and GM have low accuracy due to low specificity and sensitivity, respectively [24]. Machine learning-based approaches such as the random forests classifier appear to perform much better than AC an GM [10, 15].

#### Uni- and Bimanual upper-limb use

The upper-limb use signals from the two limbs can be used for defining uni- and bimanual upper-limb use at time *t* as the following:

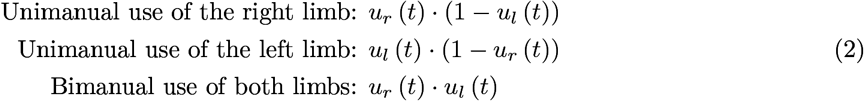

### 2.3 Instantaneous Intensity of Use (IIU)

*Definition* **Instantaneous Intensity of Use** (IIU) is a construct that reflects how strenuous a movement/posture is at a particular instance of time, when the upper-limb is in use.

Some examples of measures (*f*_*μ*_) to quantify IIU include the magnitude of movement velocity, acceleration, interaction force, muscle activity, etc. Let *μ*_*i*_ represent the IIU signal for the upper-limb *i*. It assumes non-negative values when the upper-limb is used, and is defined to be zero otherwise.

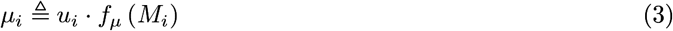

where, *μ*_*i*_ ∈ ℝ_*≥*0_([0*, T*]), and the function 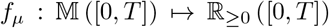 computes IIU signal from the upper-limb measurement signal *M*_*i*_.

The exact choice for *f*_*μ*_ is application-specific and dictated by 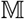. We also note that results obtained from different types of measurements and different measures *f*_*μ*_ might not be comparable, e.g. the magnitude of movement velocity can be independent of the magnitude of movement acceleration. Thus, it is imperative to report the exact *f*_*μ*_ and its units when reporting the instantaneous intensity of use. Activity counting, as defined in [8, 9, 11], is an example of an IIU measure in the current literature.

In general, *μ*_*i*_ (*t*) will not be uniformly zero in a continuous interval of time *t* ∈ [*t*_1_*, t*_2_] where there is movement (change in limb position or configuration). However, *μ*_*i*_ (*t*) can be uniformly zero in a continuous interval under two circumstances:

1. *u*_*i*_ (*t*) = 0*, ∀t ∈* [*t*_1_*, t*_2_]: When there is **no upper-limb use** during this interval.
2. *f*_*μ*_ (*M*_*i*_) (*t*) = 0*, t* ∈ [*t*_1_*, t*_2_]: When the **upper-limb is used in a meaningful posture**, *f*_*μ*_ (*M*_*i*_) (*t*) can be uniformly zero in the interval *t* ∈ [*t*_1_*, t*_2_] for some choice of measurement signal and *f*_*μ*_. For example, activity count, magnitude of movement acceleration/velocity will be zero during an upper-limb posture. On the other hand, the magnitude of muscle activity controlling the upper-limb will not be zero even while holding a voluntary posture.

It is important to be aware of these issues when interpreting time intervals where *μ*_*i*_ (*t*) is uniformly zero.

### 2.4 Windowed Upper-limb Use (WUU)

*Definition* **Windowed upper-limb use** (WUU) is a construct that reflects the proportion of time an upper-limb is used in a given time period *D*.

WUU at time *t*, denoted by 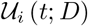, can be computed as the average value of *u*_*i*_ in the past *D* seconds.

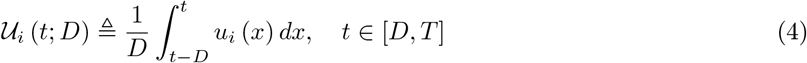

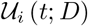 is a smoothed version of *u*_*i*_. We will drop *D* in the parenthesis in the rest of the manuscript and use it only if its explicit mention is required. From Eq. 4, we can immediately identify some essential properties of 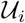:

- 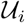 is a continuous-valued signal that can take on any value in the closed interval [0, 1].
- The value of 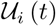 indicates the proportion of time in the interval (*t − D, t*] where the upper-limb was used, i.e. *u*_*i*_ (*t*) was 1. Thus, there are infinitely many *u*_*i*_s that can result in the same 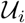.
- The value of the parameter *D* will depend on the application, and controls the amount of smoothing of *u*_*i*_; larger values of *D* will results in smoother 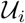 while compromising time localization of the information conveyed by 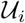. When *D* = *T*, then 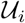 measures the proportion of time the upper-limb *i* was used over the entire measurement epoch.

### 2.5 Windowed Intensity of Use (WIU)

*Definition* **Windowed Intensity of Use** (WIU) is a construct that reflects the average intensity of upper-limb use in a given time period *D*.

WIU 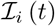 can be computed from measures of upper-limb use *u*_*i*_ and measurement signals *M*_*i*_(*t*) as the following,

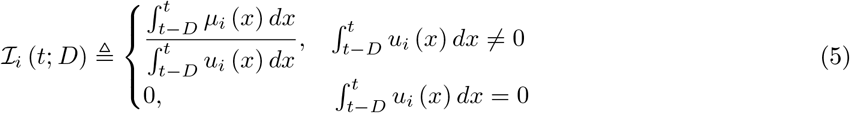

where, 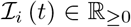. The same ambiguity as *μ*_*i*_ (*t*) exists when 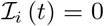 for some time *t*. 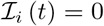 could mean that the upper-limb was either not used during the time interval (*t − D, t*] or it was used for performing upper-limb postures.

### 2.6 Windowed Upper-limb Activity (WUA)

The amount of use of an upper-limb during a measurement epoch depends on both the duration and intensity of movements performed during this period, which are captured by 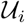 and 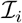, respectively.

*Definition* **Windowed upper-limb activity** (WUA) is a construct that reflects of how long and how intensely an upper-limb is used in a given time period *D*.

High amounts of average upper-limb activity correspond to long duration, high intensity movements, while low activity corresponds to short duration, low intensity movements. Windowed upper-limb activity 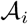 of the upper-limb *i* can be captured by the product of 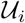 and 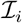, which quantifies the co-variation of these two factors. We thus define 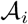 as,

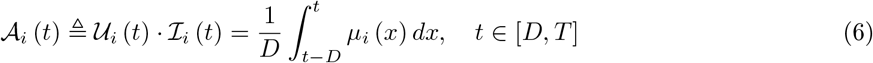

where, 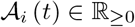 assumes non-negative values and is upper-bound by 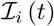. A subject with high values for 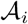 would be referred to as more active, than one with lower values of 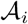. Visualization of how much an upper-limb is used during a measurement epoch, and its quantification through a single number using 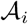 are discussed in Section 3.2.

### 2.7 Task

The four constructs – *u*_*i*_, 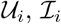, and 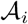 – are task-agnostic constructs that only depend on whether or not a meaningful movement or posture is performed, irrespective of its type (e.g., reaching, manipulation, drawing). To elucidate the nature of upper-limb use, task-specific measures are required, i.e. measures that can classify the types of tasks being performed, how well these tasks performed, etc. This information could be used to target therapy to accomplish specific rehabilitation goals. To carry out task-specific analysis, one must first define a set of tasks of interest that can be identified from the measurements *M*_*i*_.

*Definition* **Task** is any upper-limb movement or postural pattern of interest.

Let the set 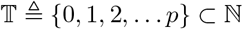 be a set of natural numbers representing the *p* distinct tasks of interest; the numbers from 1 to *p* correspond to the *p* tasks, and 0 represents all tasks other than these *p* tasks of interest. Let 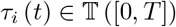 represent the task performed by the upper-limb *i* at time *t*.

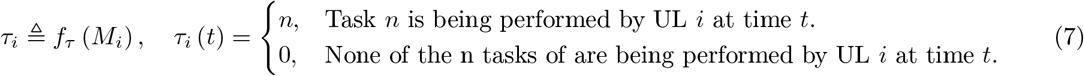

The function *f*_*τ*_ is a measure that maps the measurement signal *M*_*i*_ to *τ*_*i*_, i.e. 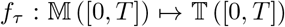. We assume that, in general, the *p* tasks of interest are functional in nature, which implies *τ*_*i*_ (*t*) can take on a non-zero value only if *u*_*i*_ (*t*) = 1. The choice of *f*_*τ*_ will depend on 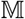 and 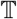. Similar to upper-limb use, the detection of tasks from the measurement data will also be probabilistic in nature due to the natural intra- and inter-subject movement variability.

### 2.8 Movement Quality (MQ)

*Definition* **Movement Quality** (MQ) is construct that reflects the quality of the underlying sensorimotor control.

MQ is a high level construct that can be expressed in terms of other constructs such as movement smoothness [18], coordination [25, 26], etc. The components of movement quality include both tasks-specific and task-agnostic constructs, which need to be computed slightly differently. We note that it might also be of interest to evaluate the quality of postures, in which case this construct could be generalized to mean movement or posture quality.

Task-agnostic measures of movement quality could be, e.g., amount of tremor, which could be computed from the *M*_*i*_ without worrying about the underlying tasks being performed. However, task-specific measures such as the ones to quantify smoothness, coordination, etc. must be computed only from complete data segments corresponding to a particular occurrence of a specific task. This is because the appropriate interpretation of such task-specific MQ measures requires the necessary contextual information, which must include at least the task being performed. There is currently little work on classifying tasks and estimating movement quality using sensors for upper-limb assessment in daily life.

## 3 Visualization of Upper-limb Functioning

Measuring upper-limb movements during daily-life can result in vast amounts of data, which need to be summarized through appropriate quantitative and graphical means. A well-designed graphical summary can provide quick and clear insights into data, and allow users to answer specific questions about upper-limb behavior. In this section, we present three graphical approaches for summarizing different types of behavioral information: (a) temporal profile of upper-limb functioning; (b) summary of upper-limb activity; and (c) relative use of the two upper-limbs. All data presented in this section were obtained from a previous study by David et. al [10]. The measurements were obtained from IMUs donned on each wrist, i.e. 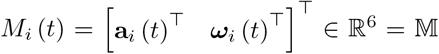 and consists of the linear acceleration **a**_*i*_ (*t*) and angular velocity ***ω***_*i*_ (*t*) measured by the triaxial accelerometer and gyroscope, respectively, at time *t* from the upper-limb *i*. Upper-limb use was estimated using the GM score algorithm [13], and instantaneous intensity of upper-limb use was chosen to be the activity counts [27] derived from the accelerometer data. Windowed upper-limb use and intensity were computed using *D* = 60*s*.

### 3.1 Temporal profile of upper-limb functioning

The plot of upper-limb use *u*_*i*_ (*t*) over the course of several measurement epoch, allows the user to see changes in upper-limb use over time. The outcomes from measures of upper-limb use could be presented in chronological order so that a clinician can see variations in upper-limb use over the course of the day or days. Sample plots of *u*_*l*_, 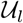, *μ*_*l*_, 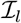 for a healthy (left column) and an impaired subject (right column) over a period of 90 minutes are shown in Fig. 3. The left upper-limb use *u*_*l*_ is visualized as an event plot in Fig. 3(a)-(b), where the presence of a vertical line at time *t* means *u*_*l*_ (*t*) = 1, else it is 0. The windowed upper-limb use 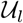 is displayed in a red trace in Fig. 3(a)-(b). Fig. 3(c)-(d) display the corresponding *μ*_*l*_ and 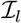 for this period in gray and blue traces, respectively; *μ*_*l*_ (*t*) = 0 whenever the upper-limb was not used or there was a functional posture. 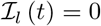 when the upper-limb was not used or used in a posture in the last *D* seconds, i.e. 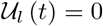.

**Figure 2:**
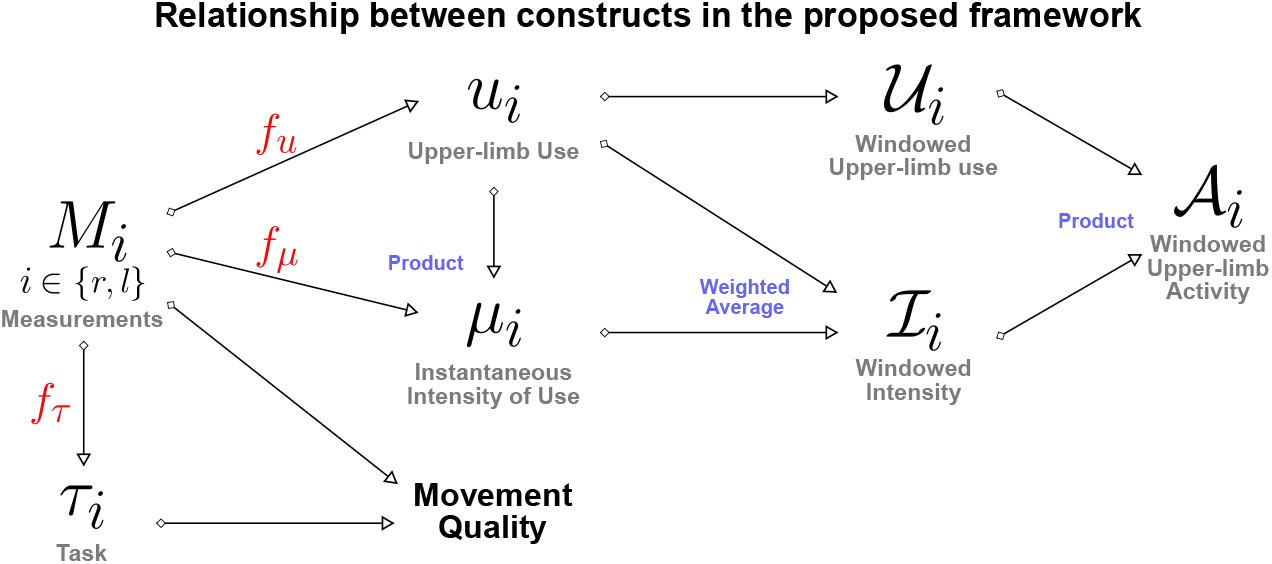
A directed graph representation of the connections between the different constructs defined in the proposed framework. The leftmost node represents the measurements, while the rest of the nodes are constructs of interest in the assessment of upper-limb functioning. The construct at the end of a directed edge is derived using the construct/measurements at the start of the directed edge. The measures (red color text) used to quantify a construct from measurements are placed above the directed edge. The blue colored text next to some of the construct indicate how two constructs are combined to derive the target construct.

**Figure 3:**
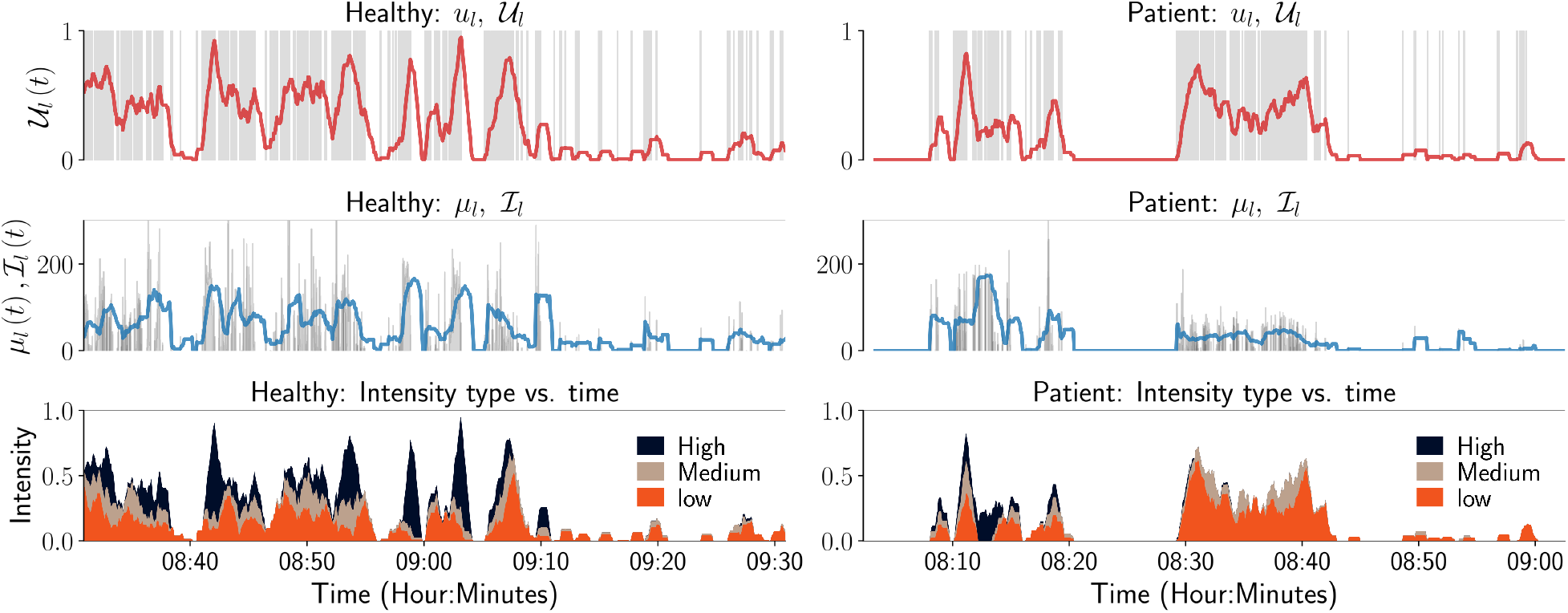
Temporal visualization of constructs related to upper-limb use and intensity. The left and right columns correspond to data from a healthy participant and a patient, respectively. The top row depicts the left upper-limb use signal *u*_*l*_ as a gray-colored event plot, where the vertical gray line at time *t* indicates *u*_*l*_ (*t*) = 1. And the light red colored graph shows the corresponding windowed upper limb use 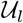. The middle row depicts the instantaneous intensity of use *μ*_*l*_ (gray) and the widowed intensity of use 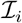 (light blue) for the left upper-limb. The bottom row depicts the proportion of time the intensity of use was low (orange), medium (brown), or high (black) in the last 60s. Although not shown in these figure, it would also be useful to indicate in such plots periods of time where there is no data available, i.e. periods where a wearable sensor has been removed and is not recording movement data from a subject.

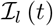 only provides a summary of the intensity of left upper-limb use in a temporal segment by computing the average intensity. A more detailed depiction of movement intensity can be provided by displaying the relative proportions of time, in an observation window, where the movement intensity is **low**, **medium**, or **high**; the definitions of the three intensity levels are provided in the figure’s caption for this particular case. The plots in Fig. 3 can aid clinical evaluation, as they indicate that the overall amount and intensity of use for the patient (right column) is lower than that of the healthy subject. The patient also has little or no high intensity movements compared to the healthy participant (Fig. 3(e) and Fig. 3(f)).

### 3.2 Visualization of upper-limb activity

A visual summary of the amount of upper-limb use during a measurement epoch can be provided through a scatter plot of 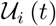 versus 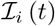, ⩝*t,* such that 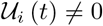^3^. This plot will be referred to as the *UVI plot*, which provides a simple visual summary to quickly gauge overall upper-limb activity. With no loss of generality, we have chosen 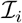 and 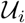 to be the *x* and *y* axes of the UVI plot, respectively. Such a plot has the following properties:

- All points of this scatter plot belong to the set *P* = {(*x, y*) | 0 ≥ *x,* 0 *< y ≤* 1}. This a strip of height 1 extending along the positive *x* axis.
- By definition, the *x* axis is not part of the plot since only data points where 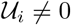 are considered.
- Depending on the measurement signal and the choice of measure *f*_*μ*_, the set of all points {(0*, y*) | 0 *< y ≤* 1} will correspond to upper-limb postures; this will not be true when 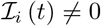 for meaningful postures.
- Scatter points with large values for *x* and low values for *y* correspond to short duration high intensity movements, e.g. swatting a fly.
- Points with values of *y* close to 1 and low values for *x* correspond to prolonged low intensity movements, e.g., writing, typing.

Data from both upper-limbs can be visualized in a single plot by plotting them in the first and second quadrants as shown in Fig. 4. Here, the right and left upper-limbs are depicted in the first and second quadrants, respectively; note that the data in the second quadrant are plotted by negating the value of 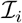. The light red colored lines in these plots correspond to constant windowed upper-limb activity lines, i.e. 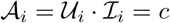, where *c* is a constant.

**Figure 4:**
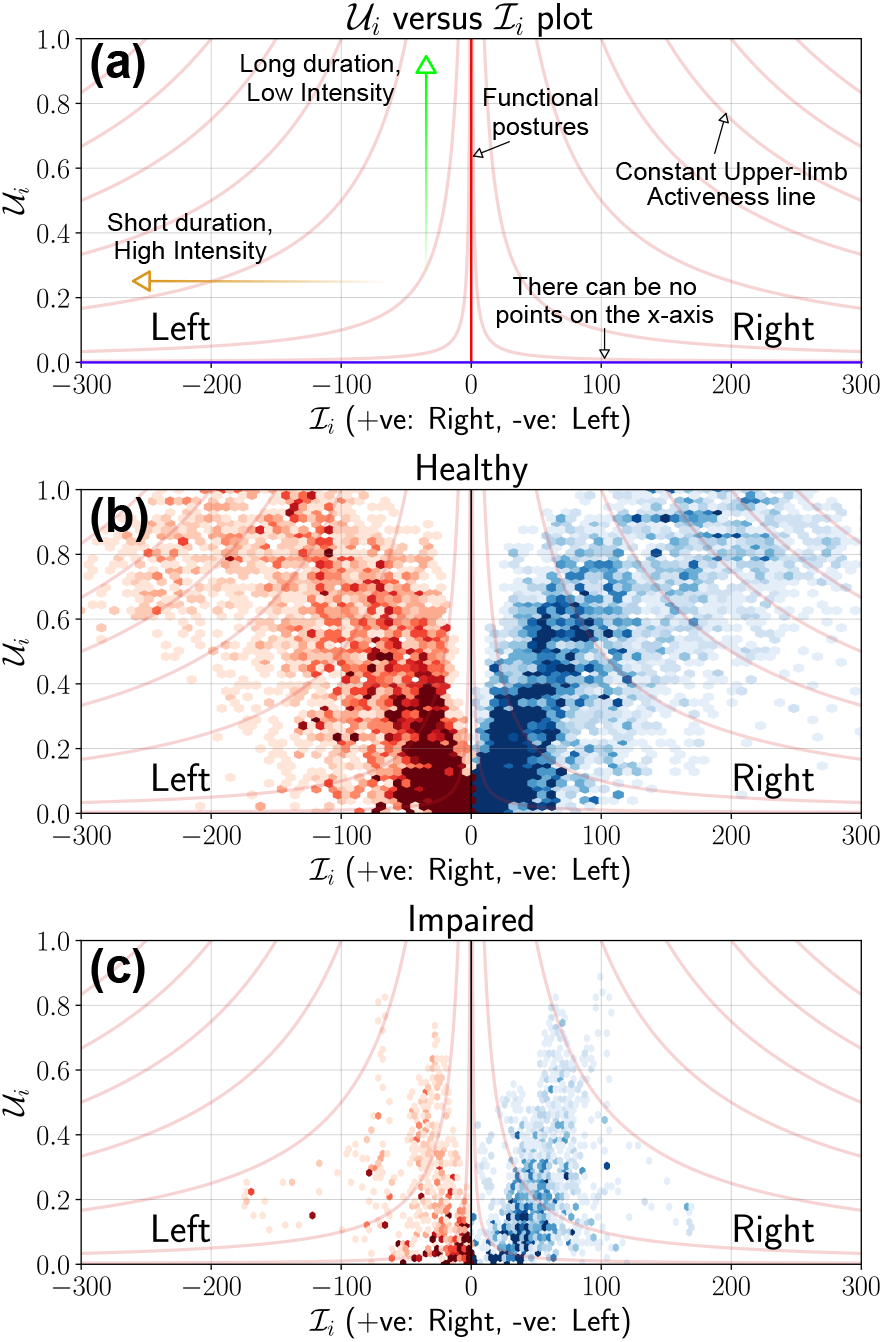
Use versus Intensity (UVI) plot to depict the overall amount of use of the upper-limbs. a) This plot provides the details of a typical UVI plot and highlights some critical elements to help interpretation. The *x* axis cannot be part of the plot, and light red colored curves are the constant upper-limb activity lines. If *f*_*μ*_ is the magnitude of acceleration as is the case in (b) and (c), then the *y* axis represents meaningful/functional postures where the intensity can be zero. (b) UVI plot for a healthy participant using data collected from a single day. The 1^*st*^ and 2^*nd*^ quadrants of the scatter plot depicts the right (blue) and left (red) upper-limbs, respectively. (c) UVI plot for a stroke participant using data collected from a single day. It is clear that the stroke participant has a low level of activity compared to the healthy participant.

Fig. 4(b) and Fig. 4(c) display the UVI scatter plots for a healthy and stroke participant, respectively, using data collected from a single day (6 to 8 hours) of recording [10]. For the healthy subject, most points are of short to medium duration 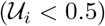 and low intensity 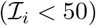 in Fig. 4(b), with some long duration, high intensity movements performed with both limbs. In comparison, most movements of the stroke participant were of relatively shorter duration 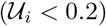, with low to medium intensity movements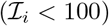; high intensity movements 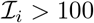 were rare. This observation is also evidenced by the constant 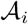 lines that cut through by the scatter plot in Fig. 4(c) compared to that of the healthy subject.

We can quantitatively summarize the UVI plot using windowed upper-limb activity 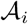 of the scatter points. The distribution of points in an UVI plot of a subject can be thought of as a sample obtained from a bi-variate probability density function of 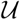 and 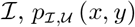^4^. The univariate probability densities of 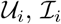, and 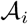 can be obtained from 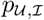 as the following,

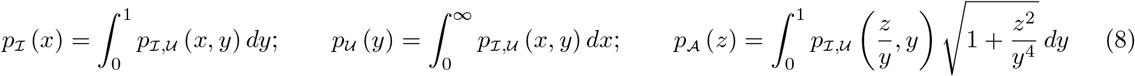

We define a quantitative measure of how much an upper-limb is used, 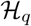, as the *q*^*th*^ percentile of 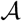, which can be computed from its probability density function 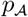,

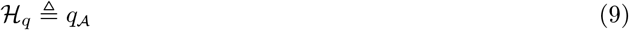

where, the subscript in *q* in 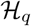 indicates that the measure is computed using the *q*^*th*^ percentile, i.e.

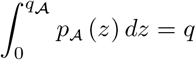

#### Properties of 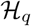

We demonstrate through a set of simulated scenarios that the measure 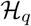 agrees with our intuition. Consider the scenarios depicted in Fig. 5, which shows five UVI plots, in the top row, with different distribution of points. In each of these plots, points are assumed to be uniformly distributed in the grey regions shown; the light red colored curves are the 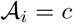 lines, where *c* is a constant. The rows of plots below the UVI plots show the univariate probability density functions 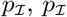, and 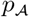 estimated from the data points sampled from the corresponding distributions 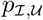 shown in the UVI plots in the top row; these plots also display the corresponding *q*^*th*^ percentile values of the sample data (*q* was set to 90).

**Figure 5:**
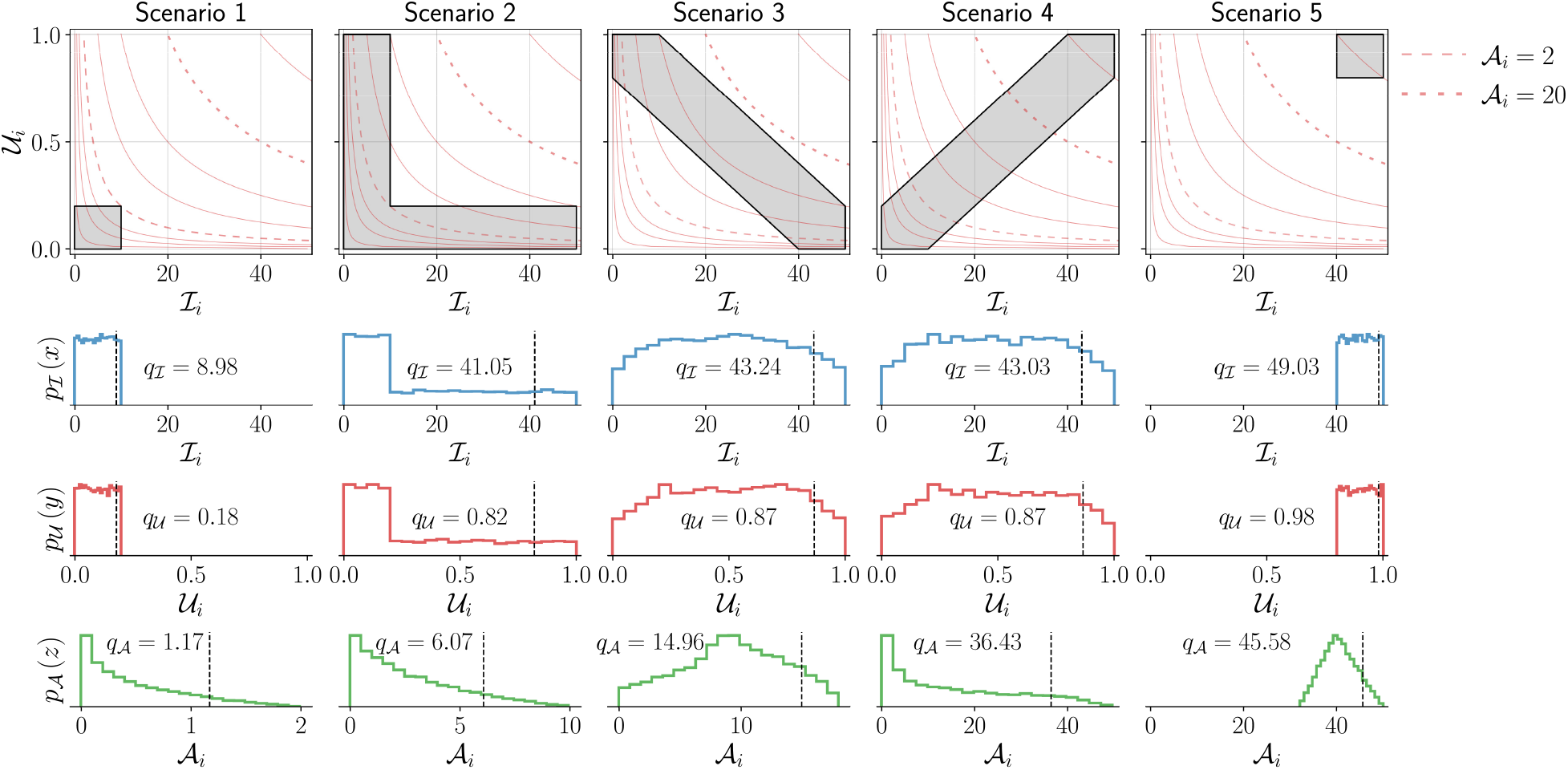
Demonstration of the measure 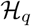 for five different simulated scenarios corresponding to different levels of upper-limb activity. The top row shows the UVI plot for the different scenarios. The shaded areas (gray) indicate different simulated scenarios from which points are sampled with uniform density. Two of constant activity lines (light red) in each plot are shown as dashed lines corresponding to 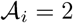 and 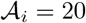. The bottom three rows depict the marginal probability density functions for 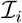 (second row), 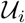 (third row), and 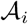 (bottom row) for these different scenarios. The black vertical dashed line indicates the *q*^*th*^ percentile (here, *q* = 90) for these different scenarios with the corresponding value written on the individual plots. Note that to enable the proper depiction of the density functions for the different scenarios, the scale for the *x* axis for the bottom row is adjusted.

The following observations can be made about the five scenarios depicted in Fig. 5, which are reflected in the measure 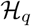:

- Scenario-1 has the lowest upper-limb activity 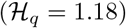 among all scenarios, as all movements are of short duration and low intensity. 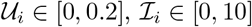, and 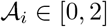.
- Scenario-5 has the highest upper-limb activity 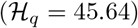 as all movements are of long duration and high intensity. 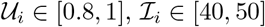, and 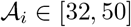.
- Scenarios 2, 3, and 4 have the same range of values for 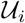 and 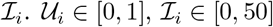.
- Scenario-2 has higher activity 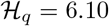 than scenario-1 as it contains movements of larger duration or higher intensity in addition to movements similar to scenario-1. This results in larger values for 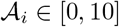 compared to scenario-1.
- Scenario-3 has higher activity 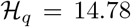 than scenario-2 as it has longer duration and higher intensity movements than scenario-2, resulting in even larger range of values for 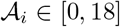 than scenario-2.
- Scenario-4 has movements with longer duration and higher intensity than scenarios 2 and 3, resulting in a large interval for the possible values of 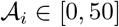 compared to scenarios 2 and 3. This results in a much higher level of activity, 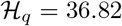.
- The difference in upper-limb activity between scenario-4 and scenario-5 is smaller than that of scenario-4 and scenario-3. Scenario-4 has more long duration and high intensity movements than scenario-3, but has more shorter duration and lower intensity movements than scenario-5. Scenario-5 only has longer duration and higher intensity movements.

### 3.3 Visualization of relative use of the upper-limbs

Visualizing the relative use of the upper-limbs has been explored through 2D scatter plots or heat-maps of different variables related to the use of the two upper-limbs [8, 10]. Relative upper-limb use can be visualized and quantified using measures of the windowed upper-limb use 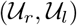, windowed upper-limb intensity 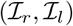 or windowed upper-limb activity 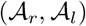; here, we use windowed upper-limb intensity for demonstration purposes. Similar to the analysis carried out with the UVI plot, we only consider the data points where at least one of the two upper-limbs was used, i.e. 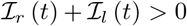^5^. It is meaningless to talk about relative use when neither of the upper-limbs are used.

In general, relative use of the upper-limbs can be visualized by plotting two functions 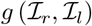 and 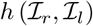 of the subject’s data along the *x* and *y* axis, respectively. These two function *g* (·) and *h* (·) will determine the nature of distribution of data points in this “*gh*” scatter plot and the plot’s fundamental properties. A qualitative understanding of these properties can be obtained by looking at the following four family of curves *L*_1_ to *L*_4_ in the *gh* plot:

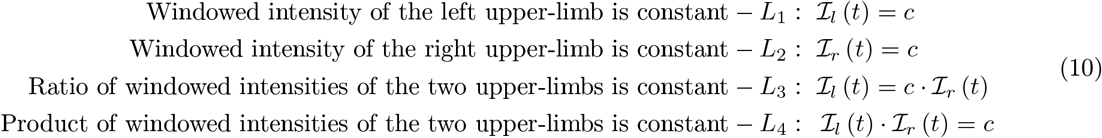

where, *c* ∈ ℝ_*≥*0_. *L*_1_ and *L*_2_ are particularly useful in explaining the shape of the distribution of points in the different visualization plots, where the bounding curves of the scatter plot are generated from different *L*_1_ and *L*_2_ curves. We present the analysis of two visualization methods, one based on the work of Bailey et. al [8] and the other from David et. al [10]:

1. Bilateral magnitude versus magnitude ratio (BMMR) plot
2. Left intensity versus right intensity (LIRI) plot

In this section we present analysis of these two visualization methods by deriving expressions for the loci of *L*_1_ to *L*_4_, and demonstrating the nature of these visualization methods using data from a healthy and hemiparetic subject [10]. We make use of the Activity Counting measure [27] to compute 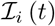. Three additional visualization methods based on BMMR and LIRI are presented in Appendix A.

#### 3.3.1 Bilateral-magnitude versus Magnitude-ratio (BMMR) plot

This method proposed by Bailey et al. [8, 9, 12] used activity counting to plot a heatmap. This is a heatmap between the *magnitude ratio* (MR) and *bilateral magnitude* (BM), which we define as the following using 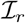 and 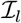,

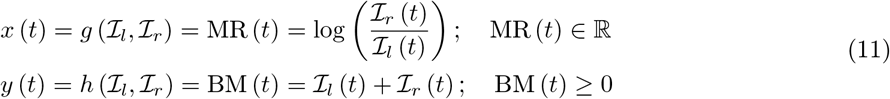

Bailey et. al bounded the value of MR to be within ±7, which we ignore in this discussion. The mathematical definitions of *L*_1_ to *L*_4_, and the plot of these curves for different values of *c* are shown in Fig. 6(a). The following are some of the essential properties of BMMR plot:

- The vertical line *x* = 0 corresponds to 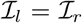, divides the plot into two halves *x >* 0 and *x <* 0 corresponding to right and left dominated halves, respectively.
- Pure unilateral use 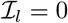 or 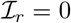 corresponds to *x* = ±∞, which was approximated to be *x* = ±7 by Bailey et. al [8].
- Equal, unbiased use of the two upper-limbs results in a symmetric leaf-like distribution of points (blue curves in Fig. 7(a) and Fig. 7(b)). The region enclosed by closed blue curve in Fig. 7(a) corresponds to 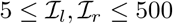.
- Biased use of the upper-limbs results in an asymmetric distribution of points, with more points located at a larger distance from the *x* axis on the side with increased use (red curve in Fig. 7(a) and Fig. 7(c)). The region enclosed by closed red curve in Fig. 7(a) corresponds to 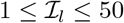 and 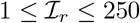.

**Figure 6:**
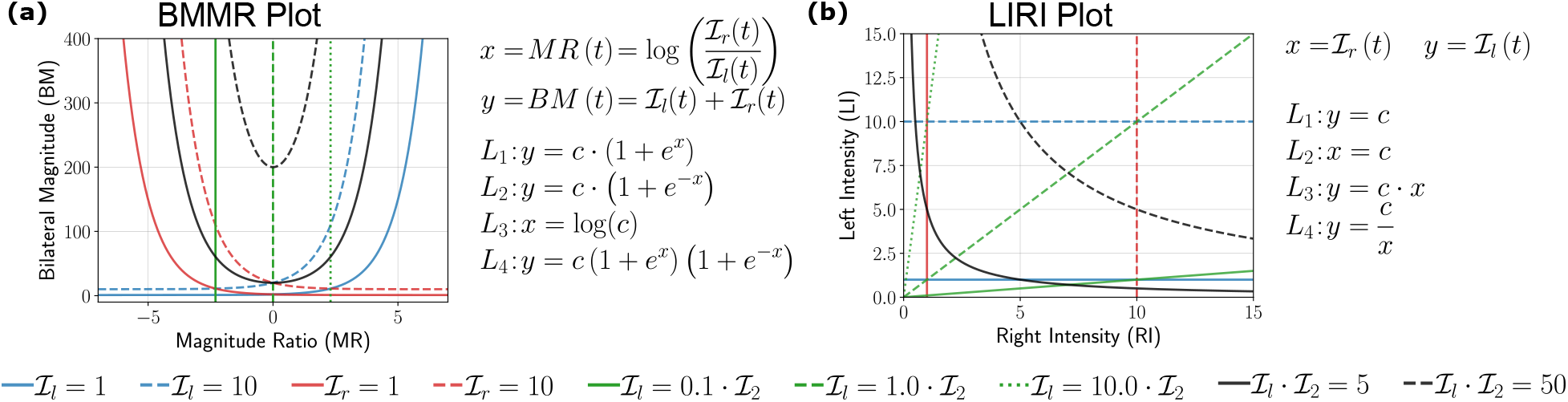
Analysis of (a) bilateral magnitude versus magnitude ratio (BMMR) plot [8] and (b) left intensity versus right intensity (LIRI) plot [10] by investigating the nature of the family of four curves *L*_1_ (blue), *L*_2_ (red), *L*3 (green) and *L*_4_ (black) introduced in Eq. 10.The solid and dashed lines indicate different values of *c* for the same curve.

**Figure 7:**
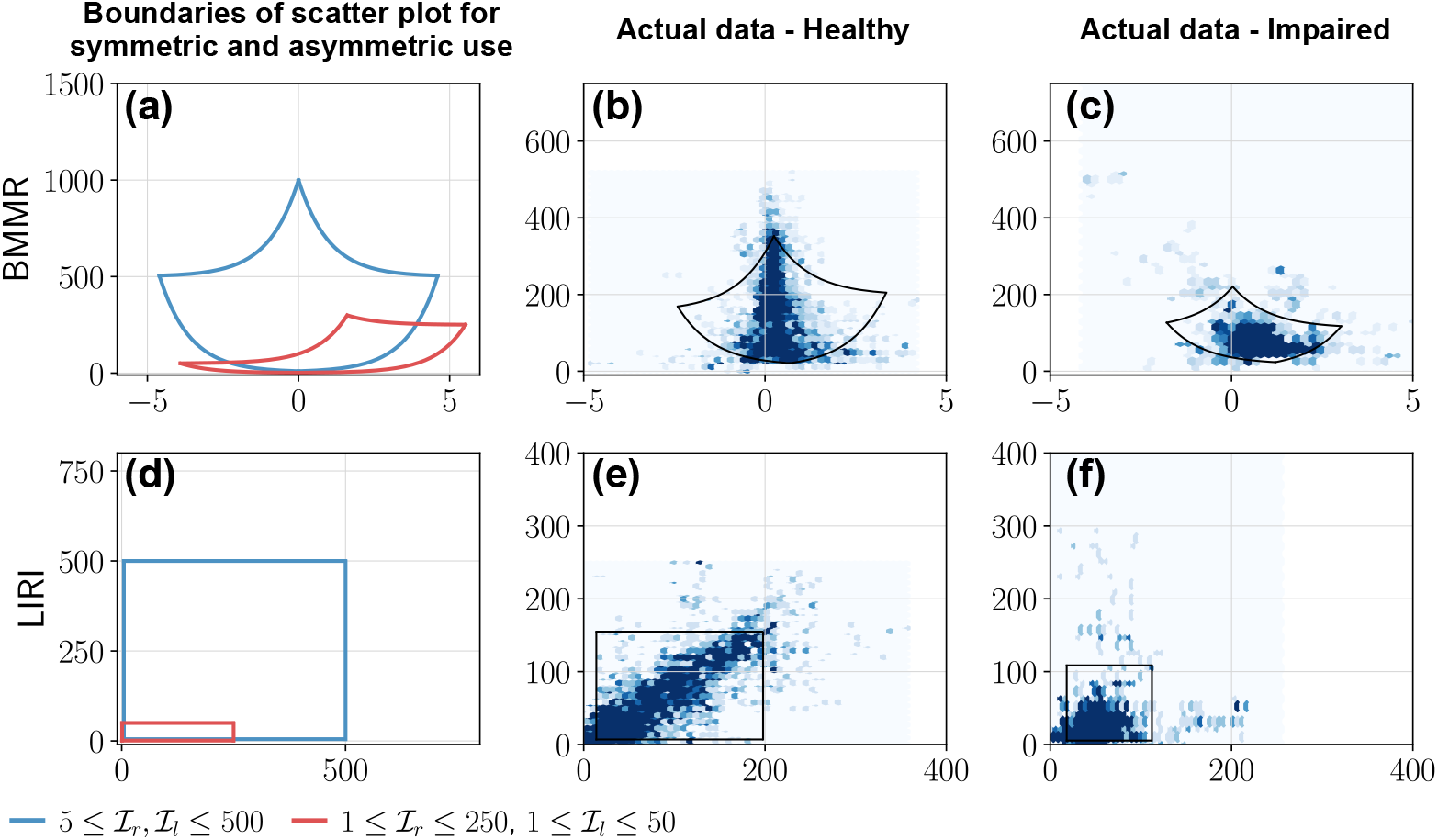
BMMR and LIRI plots of actual data from a healthy participant and a patient. The first column shows examples of the boundary of scatter plots for (a) BMMR and (b) LIRI plots for symmetric and asymmetric upper-limb use. This closed curve corresponds to the *L*_1_ and *L*_2_ curves for different values of 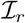 and 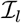. (a) The symmetric leaf shape (blue) and the asymmetric (red) shape are typical shapes seen in the plots reported by Bailey et al. [8]. Plots (b) and (c) depict the BMMR scatter plots for a healthy participant and patient using data collected during a single day. Plots (e) and (f) are the corresponding LIRI plots for the same subjects. The closed black curves shown in the plots for the healthy participant and patient correspond to the 2.5^*th*^ and 97.5^*th*^ percentiles for 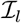 and 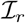.

#### 3.3.2 Left Intensity versus Right Intensity (LIRI) plot

This simple approach was proposed by David et al. [10] where the authors had used the windowed upper-limb use instead of intensity. Here, we use the windowed upper-limb intensity 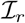 and 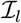 (Fig. 6(c)),

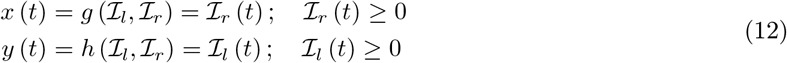

The following are some of the essential properties of LIRI plot:

- The *y* = *x* corresponds to 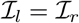 and divides the first quadrant into an upper and lower half about this diagonal line which correspond to relatively high left 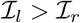 and right use 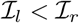, respectively.
- Pure right and left unilateral use correspond to points long the *x* and *y* axes, respectively.
- Equal, unbiased use of the two upper-limbs in a square shaped region of distribution of points (blue curve in Fig. 7(g) and Fig. 7(h)); the square is symmetric about the *y* = *x* line.
- Biased use of the upper-limbs results in rectangular distribution of points, with the longer side of the rectangular oriented along the axes corresponding to the upper-limb with increased use (red curve in Fig. 7(g) and Fig. 7(i)).

### 3.4 Quantification of Relative Upper-limb Use

A quantitative measure of relative upper-limb use should allow us to distinguish between different levels of relative use of the upper-limbs through a single number. Such a measure should map: (a) the spectrum of pure unimanual behavior to pure bimanual behavior to a compact interval on the real line, and (b) report low values for unimanual, and high values for bimanual behaviors.

We can conceive such a quantitative measure of relative upper-limb use through an approach similar to that of 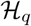. Consider the joint probability density 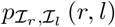 of 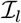 and 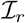^6^. We can compute the marginal densities of 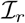 and 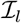, and the probability density of 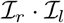 from 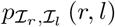 using the approach in Eq. 8. We define a measure of relative upper-limb use 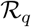 as the following,

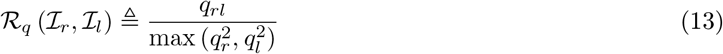

where, 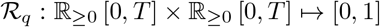 maps two time signals 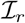 and 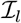 to the set [0, 1]. The subscript *q* in 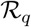 indicates that the measure is computed using the *q*^*th*^ percentiles, and *q*_*r*_, *q*_*l*_ and *q*_*rl*_ are the *q*^*th*^ percentiles of the probability density functions of 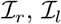, and 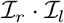, respectively. It should be noted that *q*_*r*_ and *q*_*l*_ will never be simultaneously zero as we only include data points where 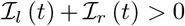. We also note that the method for computing the percentiles of the probability density functions might result in different values; this requires further investigation.

The mapping of different movement behaviors to the interval [0, 1] by this measure is shown in Fig. 8, where the LIRI plot was chosen for depicting different types of unimanual and bimanual movement behaviors. The distribution of points in these LIRI plots are indicated by the grey regions, where we assume the points are distributed with uniform density; plots with just a black line depict scenarios where the points are distributed uniformly along the line. The red diagonal line in each of these LIRI plots is the *x* = *y* line. The value of 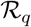 for each of these plots is shown in the respective plots, and their location in the interval [0, 1] (thick black line) on the real-line is shown in the bottom of the figure with colored vertical lines. The 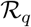 measure has the following properties.

- **Pure unimanual use**. 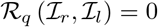 indicates pure unilateral use, such that 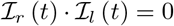, ⩝*t*^7^.
- **Symmetric bimanual use**. 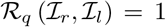 indicates pure symmetric bimanual use, such that 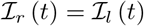, ⩝*t*.
- **Symmetry about the** *x* = *y* **line**. 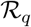 is symmetric about the *x* = *y* line, i.e., 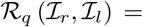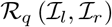. Two distribution of points that are mirror symmetric to each other about the *x* = *y* line will have the same value for 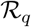. Thus, low values for 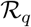 only indicate biased use and do not provide any information about the direction of the bias.
- **Bimanual asymmetric upper-limb use**. 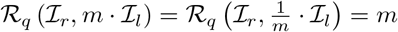, 0 *≤ m ≤* 1.
- 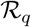 is independent of uniform scaling 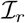 and 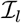, i.e. 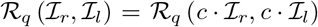, *c >* 0 is the value.

**Figure 8:**
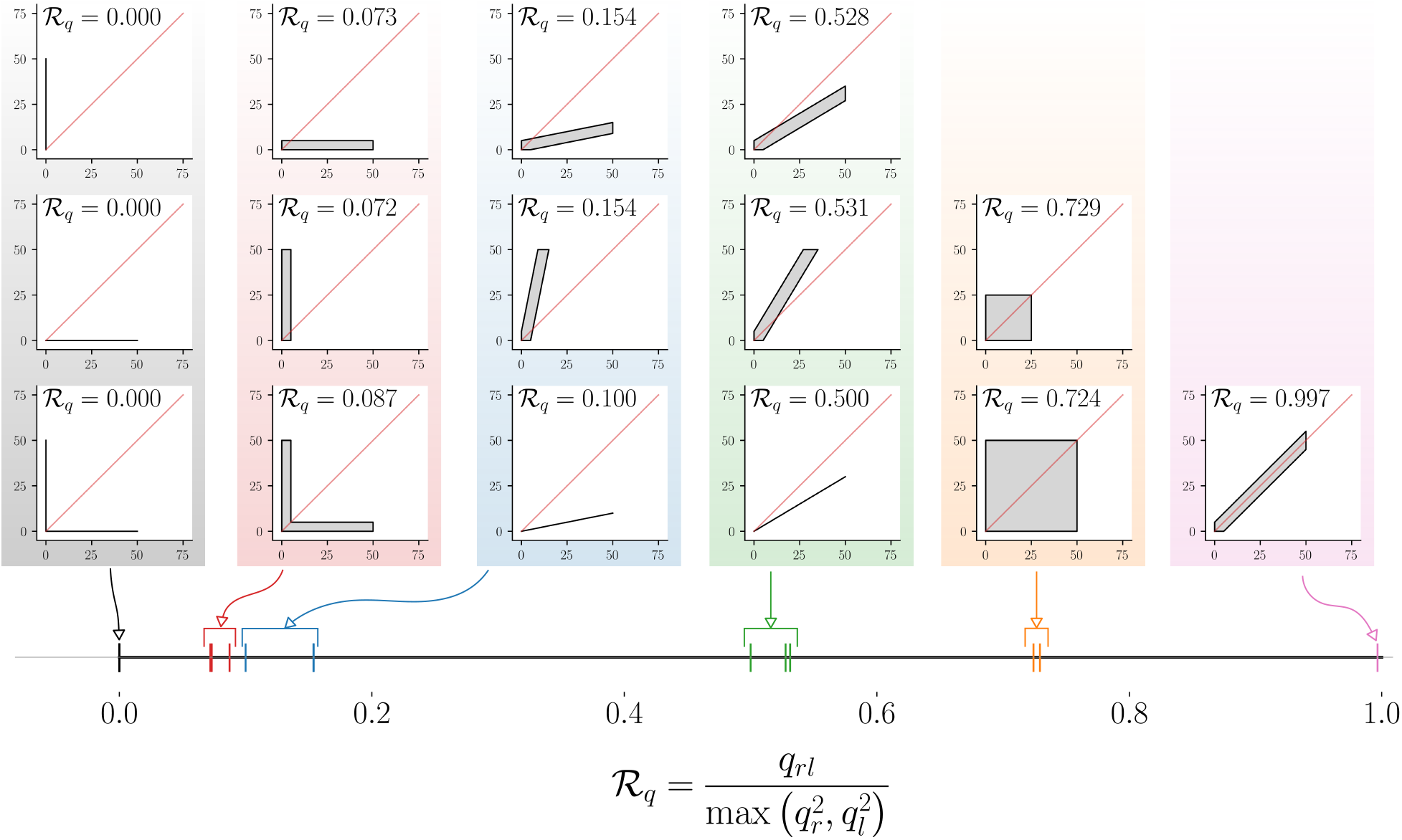
Demonstration of the mapping of different types of relative upper-limb use to 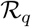. The different types of relative upper-limb use are depicted as LIRI plots grouped together to into different levels of relative upper-limb use. The leftmost column of three LIRI plots correspond to pure unimanual use. The different groups of LIRI plots from left to right correspond to reduced bias in using one limb over the other. The corresponding 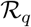 value for these different scenarios are displayed in the individual LIRI plots, and their mapping to the continuous interval [0, 1] is shown in the bottom.

The measure 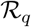 only tells us if one limb is preferred over the other, and is silent about which of the two limbs is preferred. This information can be obtained from the sign of the different between *q*_*r*_ and *q*_*l*_, which is +1 when the right limb is used more than the left, and *−*1 when it is vice versa. 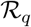 along with the sign of *q*_*r*_ − *q*_*l*_ will provide information amount of bias in using the upper-limbs, along with the preferred limb.

## 4 Discussion

The framework presented here is a step towards a rigorous foundation for sensor-based upper-limb functioning assessment by formalizing existing ideas/concepts. Lack of rigor is not an uncommon problem in movement sciences, which is reflected in the literature as ambiguous definitions of constructs, lack of clear specifications for measures, and absence of theoretical and experimental validation of measures proposed to quantify constructs of interest. Movement smoothness is a prime example of such a construct that was quantified using several measures with little or no knowledge about their properties [18, 28]. Given the increasing interest in assessment of upper-limb behavior using sensors, we strongly believe that the proposed framework can help guide future developments in this area.

### 4.1 On the importance of measurements and measures

Measurements and measures form the basis of any assessment procedure. Measurements contain “raw” information about a behavior, and measures map measurements to numbers that quantify and summarize constructs of interest. Thus, the choice of measurements and measures determine the quality of information obtained from an assessment procedure. The proposed general framework did not focus or advocate any specific measurement or measure to quantify the seven constructs (Fig. 2) defined in the framework. The two most important constructs in this framework are the upper-limb use *u*_*i*_ and instantaneous intensity of use *μ*_*i*_, which form the basis for the three constructs 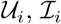, and 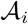 that convey information about how much the upper-limbs are used and their relative preference.

#### Upper-limb use

*u*_*i*_ is computed from the measurement signal *M*_*i*_ using a measure *f*_*u*_, both of which have implications on the accuracy of upper-limb use detection. Wrist-worn IMUs are the most commonly used sensing approach for measuring upper-limb use as these are compact, economical, and almost ubiquitous in today’s world. Although, these are ideal for picking up upper-limb movements, they are poor at detecting postures. For instance, commonly used measures that can be used to assess upper-limb use – activity counting [27] and the gross movement score [10] – cannot detect upper-limb use involving postures, or those involving pure hand movements (e.g. writing, typing). Both these methods have relatively poor accuracy in detecting meaningful movements/postures due to poor specificity or sensitivity [24]. Recent work on machine learning based methods [15, 16] have demonstrated better performance in detecting upper-limb use than existing methods. Future investigation into more sophisticated methods and the availability of more data is likely to improve this performance in upper-limb use detection. Furthermore, the incorporation of additional sensing modalities (e.g. radar-on-a-chip for hand movement tracking [29], magnetic ring finger tracking [30], EMG [31]) along with IMUs can also lead to better detection of upper-limb use. However, such multi-modal sensing approaches would need to be compact, affordable, and usable on a daily basis to ensure adoption by the end-users.

#### Instantaneous intensity of use

*μ*_*i*_ provides information about how strenuousness a movement or posture is at a particular instance of time. This is a general definition and there can be a variety of measures of *μ*_*i*_. The exact measure of intensity will be dictated by the nature of measurements 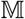. Although there are various possible measures of *μ*_*i*_, as the sensing modality for this application becomes standardized, the measure of choice for *μ*_*i*_ and other constructs are likely to converge.

### 4.2 On task-agnostic and task-specific analysis of upper-limb functioning

Upper-limb use *u*_*i*_ and instantaneous intensity of use *μ*_*i*_, and their windowed averages 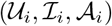 together provide a measure of how much an upper-limb is used during a measurement epoch. These constructs are independent of the nature of the task being performed by the subjects by only demarcating “functional” and “meaningiful” behaviors from non-functional ones. The work presented in the manuscript focused only on task-agnostic analysis, given these have been of primary interest in the recent literature. Although, this is necessary information, it only sheds light on the overall incorporation of upper-limbs in daily life. In particular, 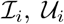, and 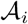 can provide information about upper-limb impairments. A more fine-grained analysis might be able to provide detailed information about specific limitations in activity and participation, which require task-specific analysis.

Task-specific fine grained analysis requires the segmentation of functional behavior into specific tasks of interest. This will allow the estimation of various impairment, activity, and participation level parameters to help build a comprehensive profile of the subject’s disability due to his/her sensorimotor condition. The details of the tasks performed during the measurement epoch provide information about limitations at the activity (e.g. time taken and range of motion while performing a task) and participation levels (e.g. restrictions in carrying out household and work-related activities). The fidelity of such detailed task-level analysis will again depend in the nature of the available measurements, and algorithms that can accurately and robustly detect the tasks of interest. There is currently no work on task-level analysis for assessing upper-limb movement behavior. This too is likely to change in the coming years with advances in human activity detection from sensors [32].

Work in the development and validation of algorithms for detecting upper-limb use and tasks are likely to benefit from the recent advances in machine learning. This work can be fast-tracked though sharing of data from various studies and groups working on this problem, since: (a) cutting edge machine learning techniques are data-hungry, and (b) collecting and collating data for validating such algorithms is time and resource intensive. The availability of such data will allow researchers with expertise in signal processing, machine learning, statistics etc. to develop algorithms to boost the accuracy of detecting upper-limb use and tasks.

### 4.3 On the visualization of upper-limb functioning

Visualization of the various aspects of upper-limb functioning is an important step in any evaluation procedure. When presenting any new visualization method, the elucidation of its fundamental properties is necessary to allow users to appropriately interpret the data presented by a graph; what constitutes a fundamental property is dictated by the information being conveyed by the proposed visualization method. These properties can often be qualitatively described through the locus of family of curves satisfying specific constraints.

The **UVI plot** proposed in this paper provides information about how much the upper-limbs are used during the measurement epoch, taking into account both the duration 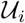 and intensity 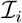 of movements. The nature of distribution of points in a UVI plot depends on several factors: (a) the nature of measurements 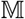; (b) the function *f*_*u*_, *f*_*μ*_ used to quantify *u*_*i*_ and *μ*)*i*; and (c) the window length *D* (Eq. 4 and Eq. 5) used to compute 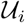 and 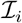. These issues were not investigated in the work presented.

Two approaches for **visualizing relative use** of the upper-limbs analysed in this paper. The family of four curves *L*_1_ to *L*_4_ (eq. 10) were used to demonstrate some properties of these visualization approaches, which helped understand the nature of distribution of points in these graphs. To promote the development and standardization of an appropriate visualization method, we make the following recommendations based on the work presented in this paper:

- **Avoiding complex transformations will make it easier to interpret graphs**. The LIRI plot is simpler than the BMMR plot, as 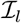 and 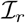 are visualized without any non-linear transformations. BMMR, MPMR, and BIUNI plots require complex transformations that hinder quick interpretation of these plots.
- **Symmetry about the** *x* = 0 **line might be easier to interpret**. Plots where the *x* = 0 line corresponds to 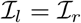 divide the plot into two regions where the use of one upper-limb is higher than the other. These plots are easier to interpret. For instance, ISID plot in appendix A, which is a rotated version of LIRI, is probably easier to interpret than LIRI.
- Any new method proposed for visualization of relative upper-limb use must be accompanied by **an analysis of at least the four family of curves** *L*_1_ **to** *L*_4_ in Eq. 10.

The visualization and quantification of relative use of the upper-limb were demonstrated using 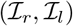. Although the properties of the visualization and quantification using 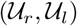 or 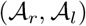 are likely to be similar, there will be some differences. One must be cautious of these differences to ensure proper interpretation of the data. For example, unlike 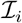 and 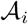 the LIRI plot with 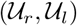 is restricted to the square 0 *≤ x, y ≤* 1.

### 4.4 Limitations

This work is an initial attempt towards a framework for the systematic analysis and interpretation of upper-limb functioning using sensors. We hope that the ideas presented here form a base for future work in this area, and anticipate that these ideas will be further refined and improved in the coming years. To aid this process, we make explicit the limitations of the current work, which are as follows:

- Most of the ideas are only presented conceptually and no algorithms or methods for quantifying the associated parameters are provided. Good algorithms for realizing the measures *f*_*u*_, *f*_*μ*_, and *f*_*τ*_ are essential for practical implementation, which will be an active area of research in the coming years.
- The components of the proposed framework are chosen based on the authors’ understanding of the current literature and the clinical needs in neurorehabilitation. The clinical usefulness of many of these ideas (concepts, measure, and visualization methods) need further validation.
- The ideas presented in the paper have been primarily targeted towards aiding the evaluation of upper-limb functioning after hemiparesis. Thus, not all of ideas presented here would be relevant for other conditions, such as those involving tremors, chorea, dystonia etc. Application of this framework to other condition, e.g. Parkinson’s disease, might require new concepts or revised definitions.
- Assessments of upper-limb functioning using sensors usually results in large amounts of data. The analysis methods that have been employed in the current literature and proposed in the current paper are quite simple, and probably only extract a portion of information available in the measured data. Future work must focus on exploring data mining algorithms for identifying patterns of recurring behavior across time. Recent developments in computational ethology [33] and automatic behavioral clustering [34] could be leveraged to identify such patterns. There is also currently little work on investigating patterns of upper-limb functioning within and across days, which might be useful in evaluating the participation of a patient in different day-to-day activities and their life roles.
- The current work only addresses questions 1 and 2 presented in the introduction section, which deal with how much the upper-limbs are used in daily life and the bias in using one limb over the other. More detailed task-level analysis are likely to be of increasing interest in the future.

## 5 Conclusion

The paper presented a framework for sensor-based assessment upper-limb functioning in a patient’s natural setting. The proposed framework provided formal definitions of essential concepts in upper-limb sensorimotor assessment, methods for visualizing assessments of upper-limb functioning, and two generic measures for quantifying the amount of upper-limbs use and the bias in using the two limbs. Demonstration of some of these components were provided through preliminary data obtained from a previous study. We also pointed out the limitations of the current work which are likely to be addressed in the coming years. We firmly believe that the work presented here will help steer future research in assessments in neurorehabilitation towards realizing an objective, accurate, and clinically relevant assessment tool to evaluate the true effect of neurorehabilitation in patients’ daily life.

## Author Contributions

AD and SB conceived the initial skeleton for the framework. The details of the framework were developed through discussions among AD, TS, VSKM, AMC and SB. AD and TS carried out the data collection and analysis presented in the paper. The initial manuscript was prepared by AD and SB. AD, TS, VSKM, AMC and SB reviewed, revised and approved the final manuscript.

## Appendix A

In this section we present analysis of three addition visualization methods by deriving expressions for the loci of *L*_1_ to *L*_4_, and demonstrate the nature of these visualization methods using data from a healthy and impaired participant [10] (similar to Fig. 7).

## Magnitude-product versus Magnitude-ratio (MPMR) plot

Instead of using the sum of *I*_*l*_ and *I*_*r*_ along the *y* axis, like in BMMR, we can use the log of the product (MP) of the *I*_*l*_ and *I*_*r*_ (Fig. 9(a)).

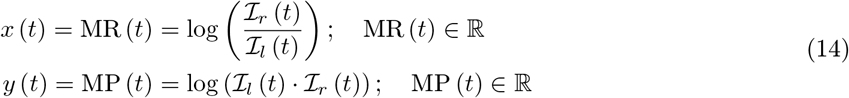

**Figure 9:**
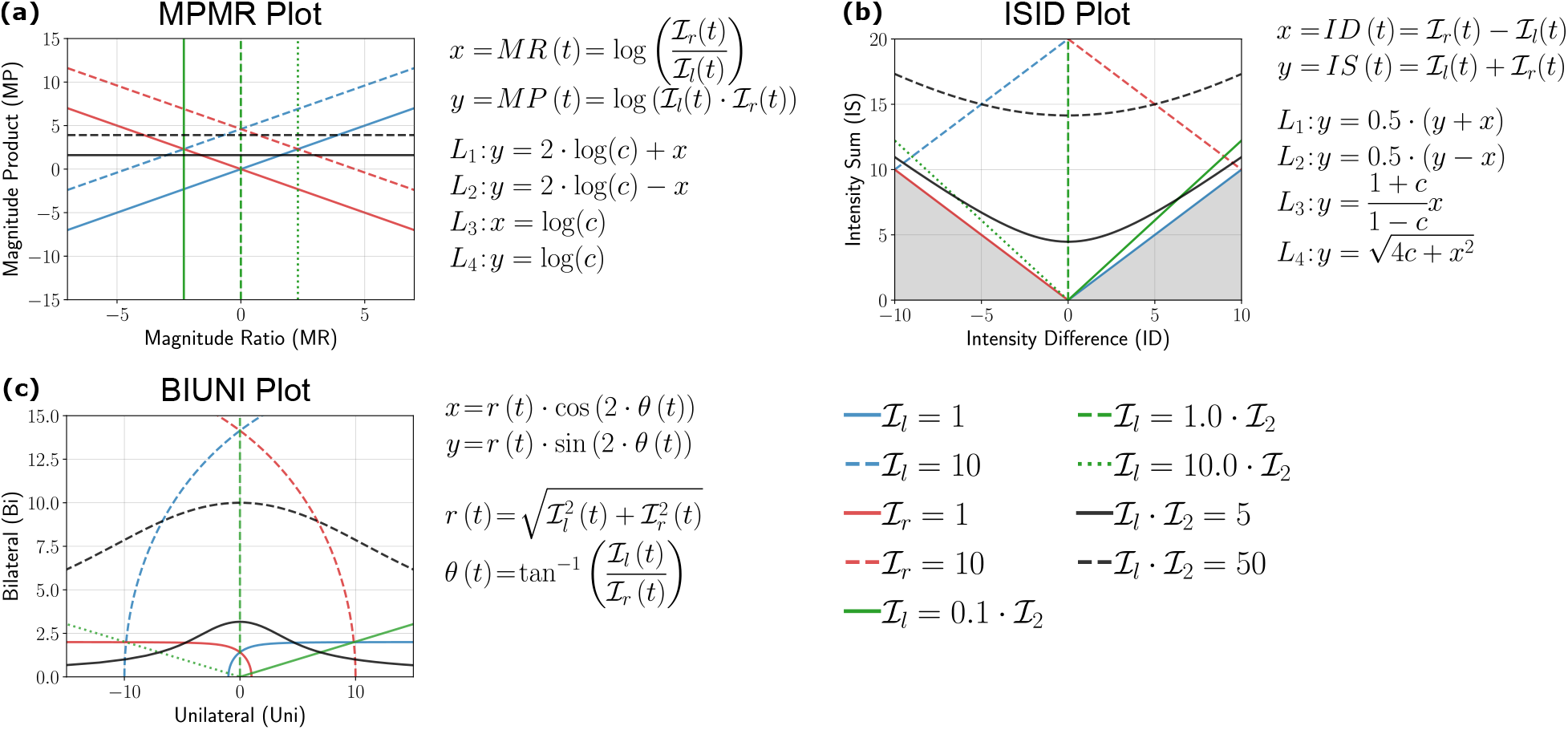
Analysis of MPMR, ISID, and BIUNI plots by investigating the nature of the family of four curves *L*_1_ to *L*_4_ introduced in Eq. 10. (a), (b), and (c) show the loci for different curves corresponding to *L*_1_ to *L*_4_ for the MPMR, ISID, and BIUNI plots, respectively.

Some of the essential properties of the MPMR plot are:

- Like BMMR, the vertical line *x* = 0 divides the plot into right and left dominated halves (*x >* 0 for right and *x <* 0 for left).
- Pure unilateral use 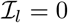 or 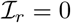 corresponds to *x* = *±∞*.
- The use of logarithmic transformation for the *x* and *y* axes leads to simpler looking curves for *L*_1_ and *L*_2_, and consequently simpler bounding regions as seen in Fig. 10(a).
- Equal, unbiased use of the upper-limbs results in a square-shaped region of distribution of points that is symmetric about the *x* = 0 line (blue curve in Fig. 10(a) and Fig. 10(b)).
- Biased use of the upper-limbs results in a rotated rectangular region of points, with more of the rectangle located on one side of the vertical line (red curve in Fig. 10(a) and Fig. 10(c)).

**Figure 10:**
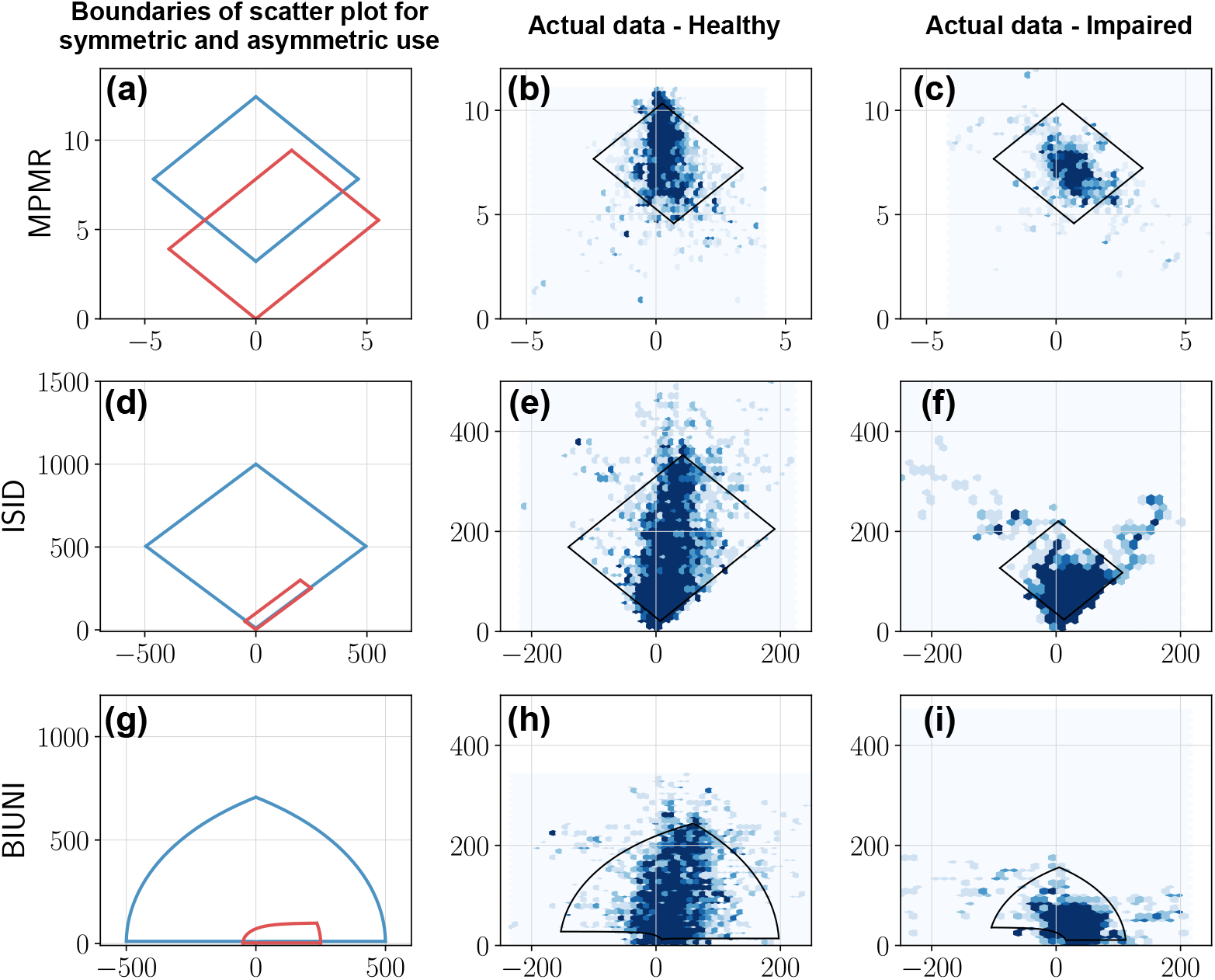
MPMR, ISID, and BIUNI plots of actual data from a healthy and impaired participant. The first column shows ((a), (d), and (g)) examples of the boundary of the distribution of scatter plots for the MPMR, ISID, and BIUNI plots for symmetric and asymmetric upper-limb use. This closed curve corresponds to the *L*_1_ and *L*_2_ curves for different values of 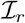 and 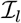. Plots (b) and (c) depict the MPMR scatter plots for a healthy and impaired participant using data collected during a single day. Plots (e) and (f) are the corresponding ISID plots for the same subjects. Plots (h) and (i) correspond to the BIUNI plots for the same subjects. The closed black curves shown in the plots for the healthy and impaired participant correspond to the 2.5^*th*^ and 97.5^*th*^ percentiles for 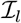 and 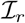.

## Intensity sum versus Intensity difference (ISID) plot

This plot is derived by rotating the LIRI plot by 45deg counter-clockwise, which results in a plot of the sum versus the difference between the average upper-limb intensities (Fig. 9(d)).

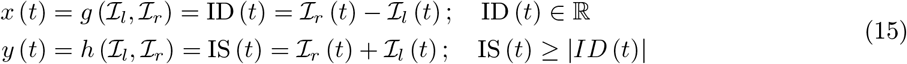

Because 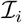 is non-negative, the points *y < |x|* are not part of the plot, which is shown by the shaded region in Fig. 9(b). The following are some of the essential properties of ISID plot:

- Like the BMMR, MPMR plots, the *x* = 0 corresponds to 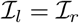.
- Pure right and left unilateral use correspond to points long the *y* = *x* and *y* = *−x* lines, respectively.
- The shape of the distribution of points are the same as LIRI but are rotated by 45deg counter-clockwise (Fig. 10(e)-(f)).

## Bimanual versus Unimanual (BIUNI) plot

This plot is obtained through a nonlinear transformation of the LIRI plot such that any point (*x, y*) in the LIRI plot with polar coordinates (*r, θ*) is mapped to a point (*r,* 2*θ*).

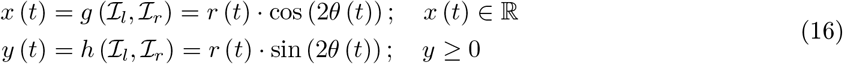

where, 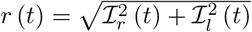 and 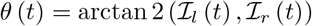.

Some of the essential properties of BIUNI plot are:

- Like the BMMR, MPMR, ISID plots, the *x* = 0 corresponds to 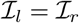.
- Pure right and left unilateral use correspond to points long the positive and negative *x* axes, respectively.
- Equal, unbiased use of the upper-limbs results in a symmetric dome-shaped region distribution of points (Fig. 10(h)). Biased use of the upper-limbs distorts this shape resulting in more points distributed along the side of increased use (Fig. 10(i)).

## Footnotes

2 IMU - Inertial Measurement Unit - consists of an accelerometer and a gyroscope.

3 This condition ensures that data segments where the upper-limb is at rest or is being used for non-functional movements are ignored.

4 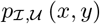 is actually a conditional density function, since we only consider data where 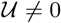.

5 As discussed earlier, 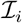 can be zero during functional postures for some measurement signals and function *f*_*μ*_. Thus, under this scenario the condition 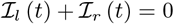 will include data either when both upper-limbs are not used or when both are used for performing functional postures.

6 This is again a conditional probability density function as we only consider data points where at least one of the upper-limbs is used, i.e. 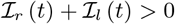.

7 It should be noted that, depending on the measurements and *f*_*μ*_, pure unimanual does not always mean the other limb is not used, since 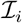 can be zero when an upper-limb is used for functional postures.

